# Pangenome Reveals Gene Content Variations and Structural Variants Contributing to Pig Characteristics

**DOI:** 10.1101/2024.10.11.617832

**Authors:** Heng Du, Yue Zhuo, Shiyu Lu, Wanying Li, Lei Zhou, Feizhou Sun, Gang Liu, Jian-Feng Liu

## Abstract

Pigs are among the most essential sources of high-quality protein in human diets. Structural variants (SVs) are a major source of genetic variants associated with diverse traits and evolutionary events. However, the current linear reference genome of pigs limits the presentation of position information for SVs. In this study, we generated a pangenome of pigs and a genome variation map of 599 deep-sequenced genomes across Eurasia. Moreover, a section-wide gene repertoire was constructed, which indicated that core genes were more evolutionarily conserved than variable genes. Subsequently, we identified 546,137 SVs, their enrichment regions, and relationships with genomic features and found significant divergence across Eurasian pigs. More importantly, the pangenome-detected SVs could complement heritability estimates and genome-wide association studies based only on single nucleotide polymorphisms. Among the SVs shaped by selection, we identified an insertion in the promoter region of the *TBX19* gene, which may be related to the development, growth, and timidity traits of Asian pigs and may affect the gene expression. Our constructed pig pangenome and the identified SVs provide rich resources for future functional genomic research on pigs.

## Introduction

As one of the most important livestock animals, pigs have a long shared history with humans and have sustained billions of people as a primary source of protein [1]. In addition to the traditional roles of pigs, a paradigm shift has occurred in recent years, with pigs emerging as an indispensable biomedical model owing to their profound anatomical, genetic, and physiological similarities to humans [2,3]. The publication of the pig genome contributed significantly to dissecting the genetic basis of distinct phenotypes and understanding the evolutionary processes of porcine domestication [2]. Although considerable advancements have been made to the Sscrofa11.1 assembly, which spans 2.5 Gb with 93 gaps, predominantly on chromosome Y, it falls short of encapsulating the full spectrum of genetic diversity across global pig breeds [4]. Despite ongoing refinement efforts, the reliance on a solitary female Duroc pig as the basis for this reference genome remains a limitation [5]. Comparative analyses of recently generated chromosome-level porcine assemblies have demonstrated substantial genetic variation among diverse breeds, particularly among SVs [6]. An increasing number of studies have indicated the pivotal roles played by SVs in genome evolution and genetic control of economic traits in domestic animals [7,8]. However, conventional linear references face challenges in detecting large SVs and accurately inferring the genotypes for each locus [9]. The pangenome concept has been proposed and iteratively refined to overcome this limitation, thus providing a transformative approach to addressing the limitations above.

Pangenome is a reference that incorporates genetic diversity across diverse populations within a species [10]. Unlike conventional references based on single individuals, the pangenome provides a holistic representation of the gene contents of a species [10]. A contemporary trend in pangenome studies involves the construction of graph-based pangenomes that offer enhanced versatility as references [11]. Graph-based pangenomes demonstrate superior efficacy in genotyping SVs using low-cost short-read sequencing technologies compared to linear reference genomes. This attribute facilitates the precise and efficient identification of genomic diversity within a species [12]. Recent advancements in pangenome research have led to the development of comprehensive references for human populations [13] and animals [14,15]. With the development of pangenome mapping, identification, and genotyping algorithms, an increasing number of graph-based pangenomes have been constructed, and their utility in elucidating the genetic basis of diverse phenotypes across breeds within a species has been demonstrated [16]. In pigs, a pangenome has been constructed through a comparative *de novo* assembly approach, which scrutinized 11 scaffold-level genomes from various pig breeds and augmented the existing Sscrofa11.1 reference with an additional 72.5 Mb of novel sequences that account for approximately 3% of the genome [17]. However, the complete gene set in pigs remains obscure, and the extent of SVs and their biological effects on specific traits remain underexplored.

Here, we present the *de novo* assembly of three chromosomal-level genomes seamlessly integrated into a comprehensive pig pangenome alongside existing assemblies. The gene content within this pangenome was delineated, which elucidated its core, softcore, dispensable, and private genomes and cladistic differences in the member populations. We comprehensively investigated SVs using this pangenome, revealing a distinct SV distribution across Eurasian pigs. In combination with heritability estimations, expression quantitative trait locus (eQTL) mapping, genome-wide association studies (GWAS), and selective sweep analyses, we explored the role of SVs in determining missing heritability, molecular basis of trait variation, and diversification of various breeds. The constructed pangenome and detected genetic variants provide valuable resources for future functional genomic studies on pigs and will accelerate variome-based breeding in swine.

## Results

### *De novo* assembly and annotation of domestic pig genomes

To ensure that the pangenome represents the comprehensive genetic diversity of domestic pigs, we collected sequence data from 599 domestic pigs (525 were sequenced in our previous study [18], and 74 were downloaded from the public database), with an average coverage depth of more than 20× (Table S1). These 599 individuals represented a worldwide distribution of domestic pigs and consisted of 46 Asian indigenous pig populations, 14 European domestic porcine breeds, two breeds distributed in America, and a hybrid breed (**Figure 1A**). After aligning these sequence reads to the Sscrofa11.1 genome, 51,858,536 single nucleotide polymorphisms (SNPs) were identified.

**Figure 1.**
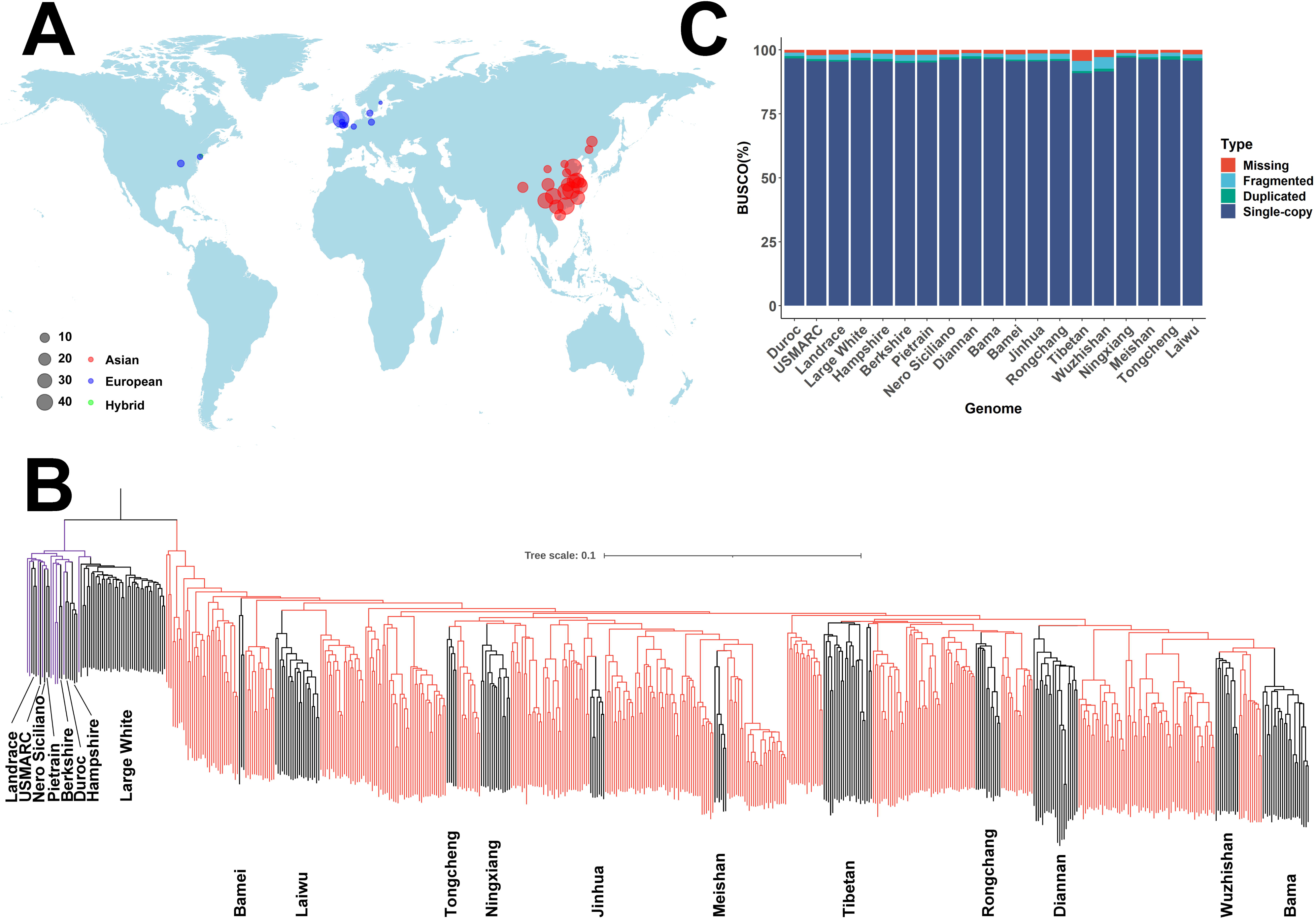
Geographic distribution and phylogenetic analysis of samples used in this study. **A.** Geographic distribution of samples used in this study. The pie size reveals the number of samples collected from each region, and the color represents the different origins (European pigs are indicated by blue, Asian pigs are indicated by red, pigs spread in America are represented by purple, and hybrid pigs are shown by yellow). The ggmap package in R generates this world map, and the map data is also achieved from this package. **B.** The whole-genome SNPs constructed a phylogenetic tree of all samples. The line with different colors indicates European pigs (blue) and Asian pigs (red). The three breeds for the new *de novo* assembly and 16 breeds whose genomes had been assembled were shown by black lines in the phylogenetic tree. **C.** The genome completenesses for 19 assemblies were assessed by BUSCO.

Based on these SNPs, the phylogenetic tree divided the 599 individuals into two large subpopulations, distinctly characterized as Asian and European pigs. Notably, according to a previous investigation, the two American and hybrid breeds were inclusively categorized as European pigs [19]. This taxonomic stratification aligns with previous findings indicating the divergence of the pig species (*Sus scrofa*) into Asian and European lineages approximately 1.2 million years ago [20]. Additionally, we discovered that Asian pigs could be classified further into four groups: BL subgroup (Bamei and Laiwu pigs), TNJM subgroup (Tongcheng, Ningxiang, Jinhua, and Meishan pigs), RT subgroup (Rongchang pig and Tibetan wild boars), and DWB subgroup (Diannan Small-ear, Wuzhishan, and Bama Xiang pigs) (Figure 1B, Figure S1). To construct a representative pangenome, 15 genomes meeting the Benchmarking Universal Single-Copy Orthologs (BUSCO) criteria (> 90% completeness; Figure 1C, Table S2) [21] were judiciously selected, thus providing a supplement to the Sscrofa11.1 genome. Three specific breeds were targeted for *de novo* assembly analysis to ensure a minimum of two assemblies per group for robust pangenome exploration.

To capture the distinct traits exhibited by Chinese domestic pigs, characterized by high reproductivity, superior meat quality, and disease resistance, we employed the PacBio Sequel II platform to sequence three characteristic breeds: Meishan pig (noted for high reproductive capacity), Laiwu pig (renowned for superior meat quality), and Tongcheng pig (recognized for disease resistance, Table S3). This effort yielded a robust dataset comprising 315.45 – 355.51 Gb of data for each individual, with an estimated coverage depth of 126 – 142× for different genomes. The assembled contig N50 values spanned from 10.93 to 17.90 Mb (Table S4). The contigs of these three genomes were corrected using corresponding deep whole-genome sequencing data. These corrected contigs were ordered, oriented, and clustered into 20 chromosomes using Hi-C data (Figures S2 and S3). Finally, we obtained chromosome-level genomes of the three breeds with a scaffold N50 of 137.13 – 140.06 Mb and genome length of 2.50 – 2.60 Gb. We further evaluated the completeness of the assembled genomes and found an average completeness of 97.13% (96.8% – 97.5%) (Figure 1C). Moreover, we validated the completeness and accuracy of the three newly assembled genomes by aligning short reads from the same breed against each genome. The alignment results revealed an average mapping ratio of 99.59% for each genome. Simultaneously, an average of 95.22% of the genomic regions across each assembly exhibited a mapping coverage of at least 10×, attesting to the high continuity and completeness of the three assembled genomes.

The three new Chinese domestic pig genomes were annotated using a strategy combining genomic, transcriptomic, and protein-derived evidence. This integrative approach yielded an average of 22,675 protein-coding gene models in the three genomes. These models exhibited an average of nine exons per gene and an average coding sequence (CDS) length of 166 bp (Table S5). The assessment of these gene models through BUSCO analysis revealed a high level of completeness, with an average of 92.5% for the 3,354 single-copy vertebrate genes (Table S6). Additionally, an average of 93.30% of the annotated genes demonstrated the potential for functional allocation to at least one of the following six databases: GO_Annotation, KEGG_Annotation, KOG_Annotation, Swiss-Prot_Annotation, TrEMBL_Annotation, and NR_Annotation (Table S7). These results revealed the high completeness of the gene annotation.

### Gene-based pangenome of pig

After acquiring the three *de novo* genomes and integrating the genomes with 15 previously downloaded genomes (excluding Nero Siciliano due to the absence of RNA-Seq data and inadequate annotation), pangenome analyses were conducted using a previously reported strategy [22]. Based on an analysis of the orthologs, all genes across the 18 pig genomes were categorized into 24,087 families. The cumulative gene sets exhibited an incremental pattern, reaching a plateau at n = 15 genomes (**Figure 2A**), indicating the representativeness of the 18 pig genomes. Among the total gene sets, 11,326 were detected in all 18 pig genomes and were defined as the core-gene families (47.02%). Additionally, 4394 families (18.24%) present in 16 – 17 genomes were identified as softcore genes, 8095 families (33.61%) detected in 2 – 15 genomes were categorized as dispensable genes, and 272 families (1.13%) identified in only one genome were classified as private gene families (Figure 2B). Corresponding to other studies [22], the percentage of core genes declined with the addition of pig genomes, although it did not reach a trough even at *n* = 18. Furthermore, the total number of core and softcore gene families accounted for more than half (65.26%) of the gene sets in the 18 genomes. In each genome, these families accounted for an average of 80.75% of the protein-coding genes (Figure 2D). However, each genome still has an average of 17.01% dispensable gene families. These observations indicated that each pig genome contained a subset of the total pig genes that were dominant in the individual gene sets and possessed a significant number of unshared genes.

**Figure 2.**
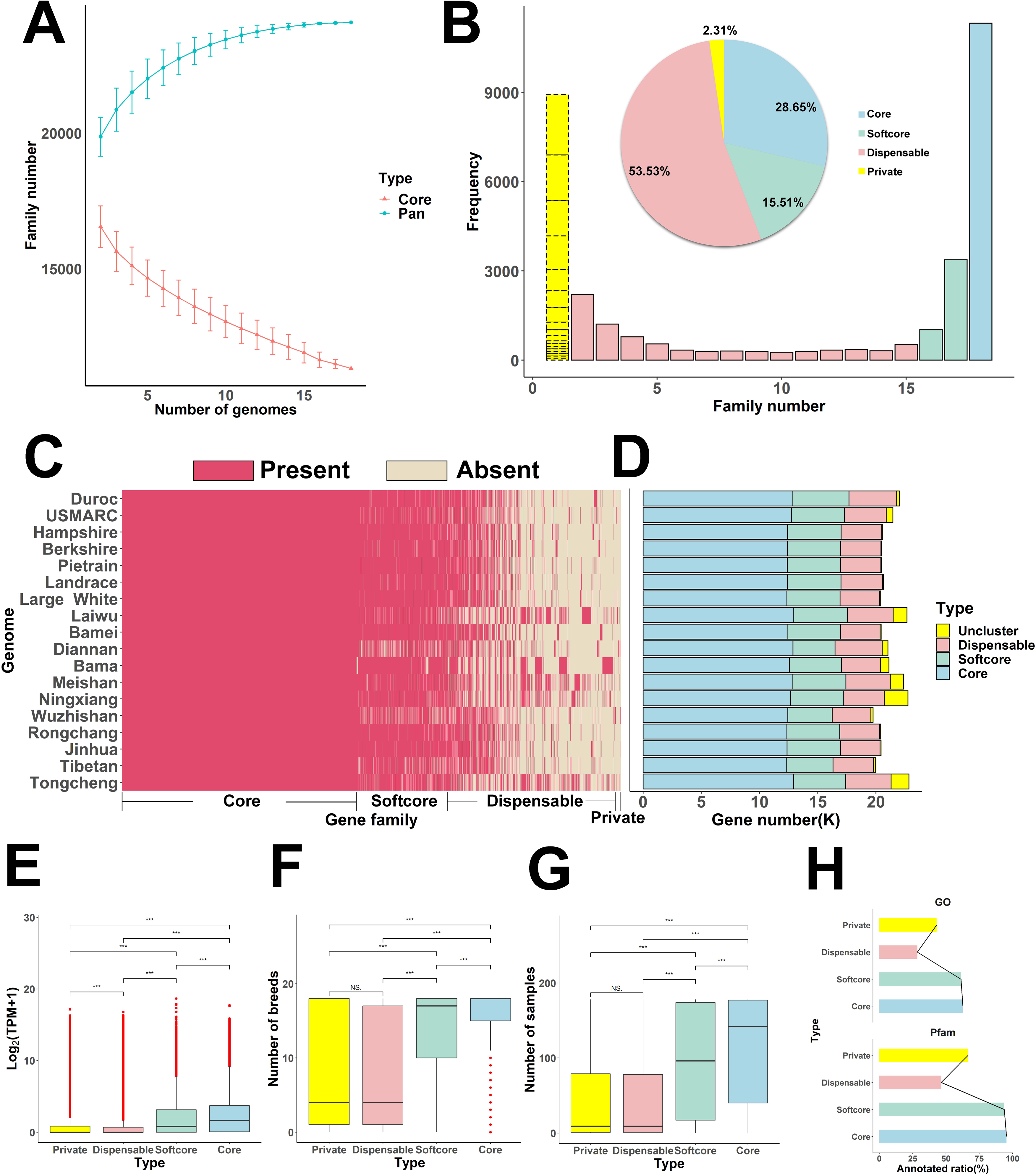
Pan– and core-genome analysis of pigs. **A**. Variation of gene families in the pan-genome and core-genome when incorporating additional genomes into the pig pangenome. **B.** Compositions of pan– and core-gene families. The histograms show the number of gene families in all genomes with different frequencies. The pie plot indicates the ratio of different gene categories. **C.** Presences and absences of gene families in each genome. **D.** Gene number of different gene categories in individual genomes. **E.** Expression profile of the pan and core genes in Sscrofa11.1. **F.** Breed-specific expression profile for pan and core genes in Sscrofa11.1. **G.** Expression width profile (indicated by expressed or not in 171 RNA-Seq data) of the pan and core genes in Sscrofa11.1. **H.** Profile of GO and Pfam annotation for the core, softcore, dispensable, and private genes.

We further investigated the expression profiles of pan genes using RNA-Seq data from 171 samples of 18 breeds (Table S8). The overall expression level, breed-specific expression status (expressed or not in the 18 breeds), and expression width (expressed or not in the 171 samples) gradually decreased from the core, softcore, dispensable, and private genes (Figures 2E, 2F, and 2G). These results suggest that the core genes exhibited a steady and broader expression spectrum, while the variable genes showed a lower, tissue-specific, breed-specific expression. Furthermore, we discovered that approximately 95.46% of the core genes and 93.83% of the softcore genes contained protein domains (reflected by Pfam), and these values were higher than the proportions of dispensable and private genes (46.52% and 66.54%, respectively). Gene Ontology (GO) annotation revealed similar results (Figure 2H), indicating that the core genes were more evolutionarily conserved than dispensable and private genes.

### Large genomic variations and hotspot regions

We aligned all 18 assemblies to the Sscrofa11.1 genome to detect genomic variants. We identified 41,724,488 SNPs in the 18 pig genomes relative to the reference Sscrofa11.1 genome (Figure S4). Although the number of SNPs from the 18 assemblies was lower than that from the 599 individuals (41,724,488 vs. 51,858,536), the distribution of SNPs from these two datasets exhibited similar patterns across the genome (**Figure 3A**). The correlation between SNP distributions from the 18 pig assemblies and 599 genomes was notably high at 0.91 (Pearson’s correlation test, P<0.05; Figure S5A). Moreover, we estimated the nucleotide diversity (π), transition/transversion ratio (Ts/Tv), and Tajima’s D value in the 18 assemblies and 599 genomes (Figure S5B, S5C, and S5D). The distribution of these features across the porcine genome exhibited high correlations between the 18 assemblies and 599 genomes, further indicating the representativeness of these 18 assemblies for Eurasian boars.

**Figure 3.**
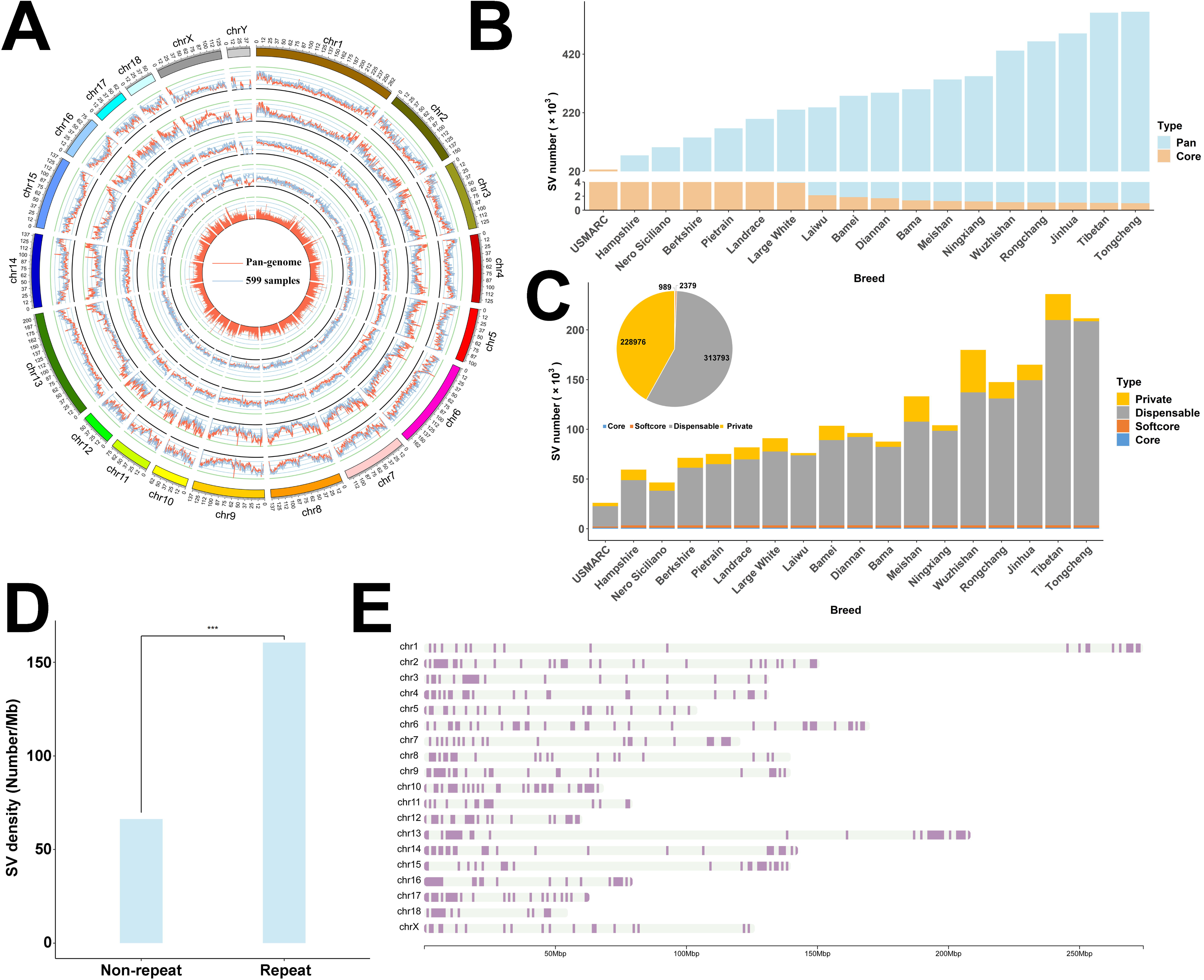
Genetic variations from 18 assemblies and 599 whole-genome sequencing individuals. **A**. Distribution of genetic variations from 18 assemblies (18 assemblies compared to the Sscrofa11.1 reference genome) and 599 whole-genome sequencing individuals. Circle plot in which concentric circles show the following (from outer to inner): ideogram of the porcine genome with colored karyotype bands, SNP density, π, Ts/Tv, Tajima’s D. The bar plot in the most inner circle represents the SV density of 18 genomes. **B.** SVs detected in each genome were merged using a non-redundant strategy, starting with USMARC and iteratively adding new SVs from additional genomes. **C.** The bar plot indicates the number of SVs for different SV categories per genome. The pie plot shows the proportion of distinct SV categories. **D.** SV density was discovered in the repeat and non-repeat genome regions. **E.** 495 SV hotspots were detected in the porcine genome.

In addition to SNPs, the assembled genomes allowed us to identify large SVs (> 50 bp in this study) by comparative genomic analysis between different breeds. We compared the 18 assemblies with the Sscrofa11.1 assembly and identified three types of SVs: presence/absence variations (PAVs), inversions, and translocations. An average of 71,704 SVs per genome were determined relative to Sscrofa11.1, and they spanned 68.70 Mb of genomic regions per breed on average. Subsequently, we merged these SVs across the 18 genomes into a set of non-redundant SVs. A total of 546,137 SVs made up of 772.92 Mb genomic regions were obtained, including 541,033 PAVs, 1887 inversions, and 3217 translocations (comprising 2231 intra-chromosome translocations and 986 inter-chromosome translocations). We discovered that the length of most PAVs ranged from 1 to 2 kb, inversions mainly ranged from 10 to 20 kb, and translocations varied from 2 to 3 kb (Figure S6). Simultaneously, the non-redundant SV set grew and tended to flatten as more assemblies were added (Figure 3B). In contrast, the set of shared SVs declined, leaving a total of 989 SVs that were shared in all assemblies.

We categorized these SVs into four classes (Figure 3C) according to their frequencies in the 18 assemblies: core (present in all 18 genomes), softcore (present in > 90% of genomes but not all), dispensable (present in more than one genome yet < 90% of genomes), and private (only detected in one genome). We found that Asian domestic pigs showed higher dispensable SVs (average of 122,106; *t*-test, P = 6.11×10^-4^) than European domestic pigs (average of 51,644, Figure 3C). However, the number of private SVs in different Asian indigenous pigs exhibited more extensive diversity (coefficient of variation [CV] = 0.87) than that in European pigs (CV = 0.34). We found that SVs tended to be enriched in repetitive DNA regions (Figure 3D). We investigated the sequence composition of each PAV. We found that 36.75% of the PAVs originated from repetitive DNA, which supports the insight that variations in repetitive sequences might contribute to the divergence of distinct genomes.

Our examination of SVs arrayed across chromosomes revealed an uneven distribution along the chromosomes, indicating that multiple and independent SVs arose in these regions. In total, we identified 495 hotspots (Figure 3E, Table S9) that overlapped with 2515 quantitative trait loci (QTLs), which were enriched in pathogen and parasite traits (P = 0.012). Furthermore, we discovered an uneven distribution of SV hotspots between distinct porcine subpopulations. In this study, we detected 2677 putative SV hotspots across five different populations (the European pig population and four distinct Asian pig populations) (Figure S7). Notably, we found 47 SV hotspots enriched over a region of 50.70 Mb on chromosome X (49.80 – 100.50 Mb) and 20 SV hotspots enriched over a region of 25.80 Mb of chromosome 5 (70.40 – 96.20 Mb) in four Asian indigenous populations; however, such enrichment was not observed in European pigs (Figure S7). In particular, the SV hotspot enrichment region on chromosome X overlapped with a previously reported region (45 – 87 Mb) and contained different haplotypes between Asian and European pigs [23]. In addition, we noticed an extended region of 42 Mb (70 – 112 Mb) on chromosome 9 that only contained an SV hotspot in European pigs. Notably, this hotspot region showed 41 independent SVs and resided in an important *CRPPA* gene that plays a critical role in skeletal muscle development, structure, and function [24].

### Impact of SVs on functional genomic regions

As SVs significantly influence genomes and are often associated with specific traits, we systematically evaluated the possible functional effects of SVs on both coding and non-coding regions. First, we discovered that most SVs were located in intergenic regions (**Figure 4A**, Figure S8). Almost half of the SVs (47.51%) were associated with at least one gene, with 43.78%, 6.83%, and 1.52% found within protein-coding genes, long non-coding RNA genes, and pseudogenes, respectively. Furthermore, we investigated the associations between the SVs identified in this study by annotating seven distinct genome features: gene, exon, intron, coding sequence (CDS), promoter, untranslated region (UTR), and enhancer. Enrichment analysis of these features revealed that most SVs were depleted in the genic areas, specifically in CDSs (Figure 4B, File S1). However, the inversion did not follow a similar depletion trend, which demonstrated enrichment across all seven features, possibly caused by the large size of most inversions. We found that 16.75% of the inversions contained an average of eight whole genes, whereas only 0.29% of the absences contained at least one whole gene.

**Figure 4.**
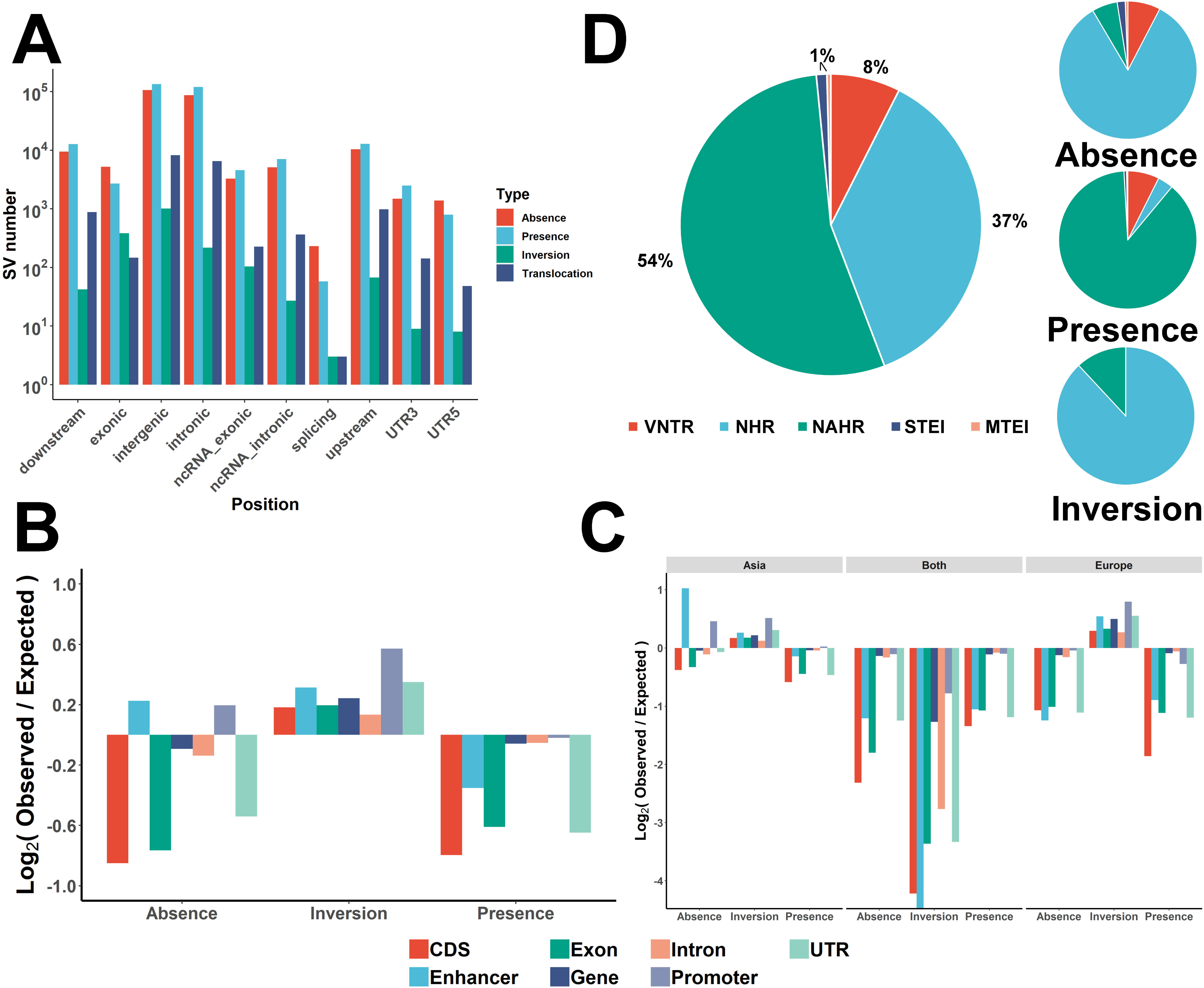
Impact of SVs on functional genomic regions and their formation mechanism. **A**. The annotation results of SVs. **B.** Enrichment/depletion of different SV types within various functional genomic features. **C.** Enrichment/depletion of different SV classes in European and Asian pigs within various functional genomic features, respectively. **D.** The proportion of SVs formed by the five estimated formation mechanisms in pigs.

Notably, absences occurred more frequently than presences in the promoter and enhancer regions. These SVs were divided into three categories: 1) SV exclusively present in European pigs; 2) SV detected only in Asian pigs; and 3) SV occurring in both groups. We conducted a comprehensive analysis to gain deeper insights into the relationship between genomic features and the three categories of SVs (Figure 4C). The results showed that the presence or absence of these three categories resulted in similar depletion signatures for genomic features, except for promoters and enhancers. In particular, the absence and presence of promoters in Asian pigs indicated enrichment of this genomic feature, which may correspond to the characteristics of Asian pigs.

### Inferring the mechanisms of SV formation

Discovering the specific mechanisms and driving forces of SV formation could improve our understanding of these large genomic variations and facilitate further studies on how large variants affect phenotypes. In this study, we employed a validated workflow [25] based on an analysis of breakpoint junction sequences of SVs to infer the mechanism of SV formation.

In total, 85.44% of the detected SVs (excluding translocations) were categorized into six types based on their inferred formation mechanisms. Among the assigned SVs, non-allelic homologous recombination (NAHR) (54.25%) was the dominant formation mechanism, followed by nonhomologous recombination (NHR) (36.74%) (Figure 4D). The observed formation percentage differed from the primary mechanism in human genomes, in which NHR is the dominant SV formation mechanism [25]. Apart from the above dominant SV formation mechanisms in pigs, our analysis indicated that approximately 44,700 SVs were shaped by variable number of tandem repeat (VNTR) and transposable element insertion (TEI, Figure 4D).

In addition, a similar investigation revealed that the dominant SV formation mechanism varied among the different types of structural variations. For instance, most presences originated from NAHR (88.09%), followed by VNTR (7.38%). However, absences and inversions exhibited distinct patterns shaped primarily by NHR. VNTR was the second most predominant formation mechanism among absences (7.64%), whereas NAHR was the second most dominant SV formation mechanism among inversions (11.92%). In addition to the SV type, we examined the dominant SV formation mechanism in different SV length categories (Figure 4D). The results showed that almost half of SVs shorter than 100 kb were formed by NAHR, while SVs longer than 100 kb were mainly formed by NHR. Notably, small SVs were more often shaped by VNTR than large SVs. In addition, concentrating on the two categories of TEI, we noticed that the size of SVs that originated from a single transposable element insertion (STEI) was mainly distributed from approximately 200 to 500 bp. In contrast, the size of SVs formed by multiple transposable element insertion (MTEI) is primarily centered at approximately 5–10 kb.

### Pangenome graph utility in pigs

We built a graph-based pangenome using Sscrofa11.1 as a linear base reference and integrated, non-redundant PAVs without many repetitive sequences (ratio < 90%). In total, 353,702 PAVs were integrated into the graph-based genome (Figure S9). We have previously reported data on 300 European-Asian hybrids [26], including genome, transcriptome, and microbiome (sampled from the feces, cecum content, ileal content, ileal mucosa, and cecum mucosa) data, which allowed us to investigate whether the pangenome graph detected SVs that lead to phenotypic variations in any traits. To test the power of the graph pangenome in capturing missing heritability, we used the LDAK [27] method to estimate the heritability of 18,189 molecular traits, including 16,037 expression traits and 2152 microbiota traits. After quality control, 286,571 SNPs and 25,933 SVs were used in the following analyses (**materials and methods**). We analyzed the contribution of these genetic variants to molecular traits individually (SNPs or SVs) and jointly (SNPs + SVs). The average heritability estimated using SVs was higher than that using SNPs (0.46 vs. 0.43; Wilcoxon rank sum test, *P*=3.42 × 10^−82^). Heritability estimation increased when both SVs and SNP categories were incorporated into the model (**Figure 5A**). The estimated heritability of genetic variants jointly was significantly higher than that of SNPs (0.56 vs. 0.43; Wilcoxon rank sum test, *P* < 10^−230^) or SVs individually (0.56 vs. 0.46; Wilcoxon rank sum test, *P* < 10^−230^). Furthermore, although both SNPs and SVs balanced the heritability of most molecular traits, traits predominantly explained by SVs still existed (Figure 5B, File S1). These results indicate that SVs identified using the pangenome approach can help capture missing heritability.

**Figure 5.**
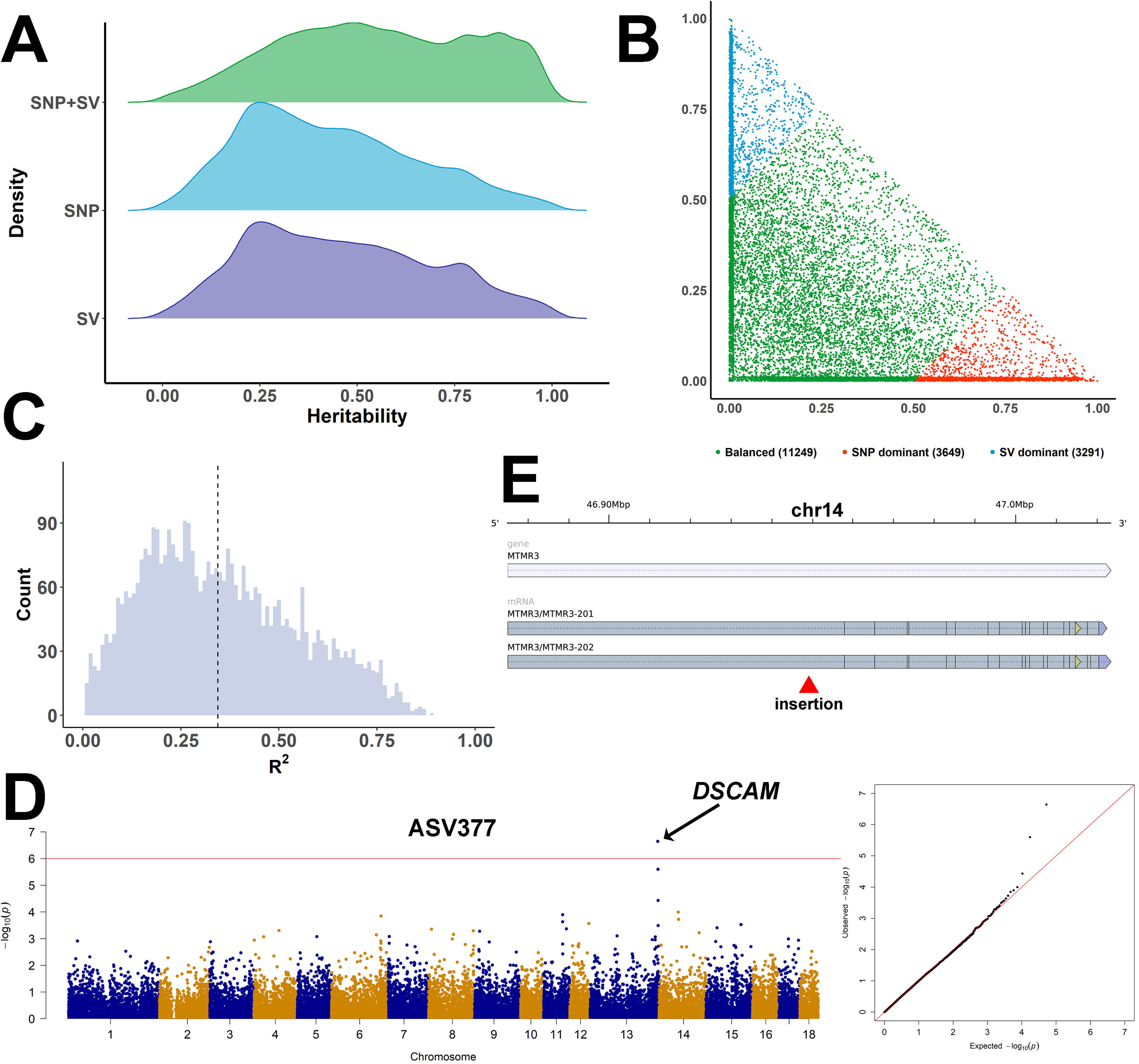
The facility of graph pangenome. **A**. Comparison of heritability estimated using different categories of genetic variants. **B.** The heritability of diverse traits contributed by SNPs or SVs. **C.** The distribution of LD between eQTLs detected by SVs and SNPs for the same genes. The dashed line represents the average of the LD values. **D.** The SV-based GWAS result for ASV337. The left is the Manhattan plot, and the right is the quantile-quantile plot. **E.** The gene structure of the *MTMR3* gene and the position of the insertion in its intron. The horizontal blue bar represents the gene region, and the horizontal grey bar indicates the transcript. The red arrow shows the breakpoint of insertion.

In addition, we detected *cis*-expression quantitative trait loci (eQTLs) based on the SNPs and SVs. Among 3137 genes, *cis*-eQTLs were detected using both SNPs and SVs. Notably, the linkage disequilibrium (LD) between *cis*-eQTLs, determined separately based on SNPs and SVs for each gene, was low (average *R*^2^ = 0.34). Only a small proportion of SVs (0.96%) were in high LD (*R*^2^>0.8) with the SNPs (Figure 5C). This result indicates low LD between different marker types is common in this public population. In addition, 139 expression genes in this population were detected in eQTLs based on SVs but not on SNPs; this could partially indicate that the SVs detected using the graph-based pangenome assisted in recognizing causal variants.

Furthermore, we performed a GWAS based on SVs to explore the correlation between SVs and the gut microbiota composition in the public population. The 16S rRNA tags (V3 – V4) clustered into amplicon sequence variants (ASVs). In the fecal samples, we identified two lead SVs associated with ASV377 and ASV2205 (Figure 5D, Figure S10). In particular, for ASV377 assigned to the *Bacteroidales* order (Table S10), we identified a tag-SV in the intron of the *DSCAM* gene associated with its abundance. This gene has been implicated in immune specificity and memory [28], and the products of this gene are particularly effective against invading parasitic or bacterial pathogens, respectively, and can even be related to gut microbiota management [29].

In addition, the graph-based pangenome provided an opportunity to detect important functional genes. In our 599 genomes, 932 tag-SNPs from a previous GWAS atlas [30] were obtained and used to search for SVs with strong LD with these tag-SNPs. In total, 102 SVs with high LD (LD > 0.8) were discovered, and 36 overlapped with the introns of 26 distinct genes (Table S11). Among these genes, the *MTMR3* gene drew our attention. Previous studies have reported an important role of *MTMR3* in the proliferation and differentiation of skeletal muscle satellite cells [31]. The insertion in the intron of the *MTMR3* gene (Figure 5E) showed high LD with the tag-SNP (NC_010456.5:g.47895001C>T) for the gestation length phenotype. This insertion may also be considered a marker for the gestation length phenotype, and the *MTMR3* gene could be an important gene for fetal development in utero [32].

### Contribution of SVs to the diversification of Eurasian pigs

Numerous genetic loci have undergone selection throughout the speciation process in European and Asian pigs, and many studies have suggested this phenomenon based on SNP data. In this study, we genotyped SVs in the 599 porcine genomes across Eurasia using our constructed graph-based genome. We calculated a validated statistical *V*_ST_ [33] to identify significant and genetically differentiated SVs within populations. A total of 2424 SVs representing the top 1% of *V*_ST_ values were identified, suggesting potential selective signatures (**Figure 6A**). Among these SVs, 784 genes overlapped, and functional enrichment analysis of these genes revealed enrichment in functions related to disease resistance, energy metabolism, and other relevant pathways (Table S12). These genes were within 1005 QTLs referencing 25 phenotypes (Table S13). Furthermore, we concentrated on genes whose CDSs were affected by these potentially selected SVs (Table S14). Finally, nine genes were detected, among which *RSAD2* plays a critical role in the immune response. An absence of 100 bp was identified, overlapping with the fifth exon of the *RSAD2* gene. Validation of this absence through polymerase chain reaction (PCR) results from various randomly selected breeds further confirmed its existence (Figure 6B). Furthermore, we aligned the RNA-Seq data of Asian and European pigs to Sscrofa11.1 and validated the existence of this absence (File S1, Figure S11). These alignment results indicated that this absence deduced a new splice junction for the *RSAD2* gene and might influence its expression.

**Figure 6.**
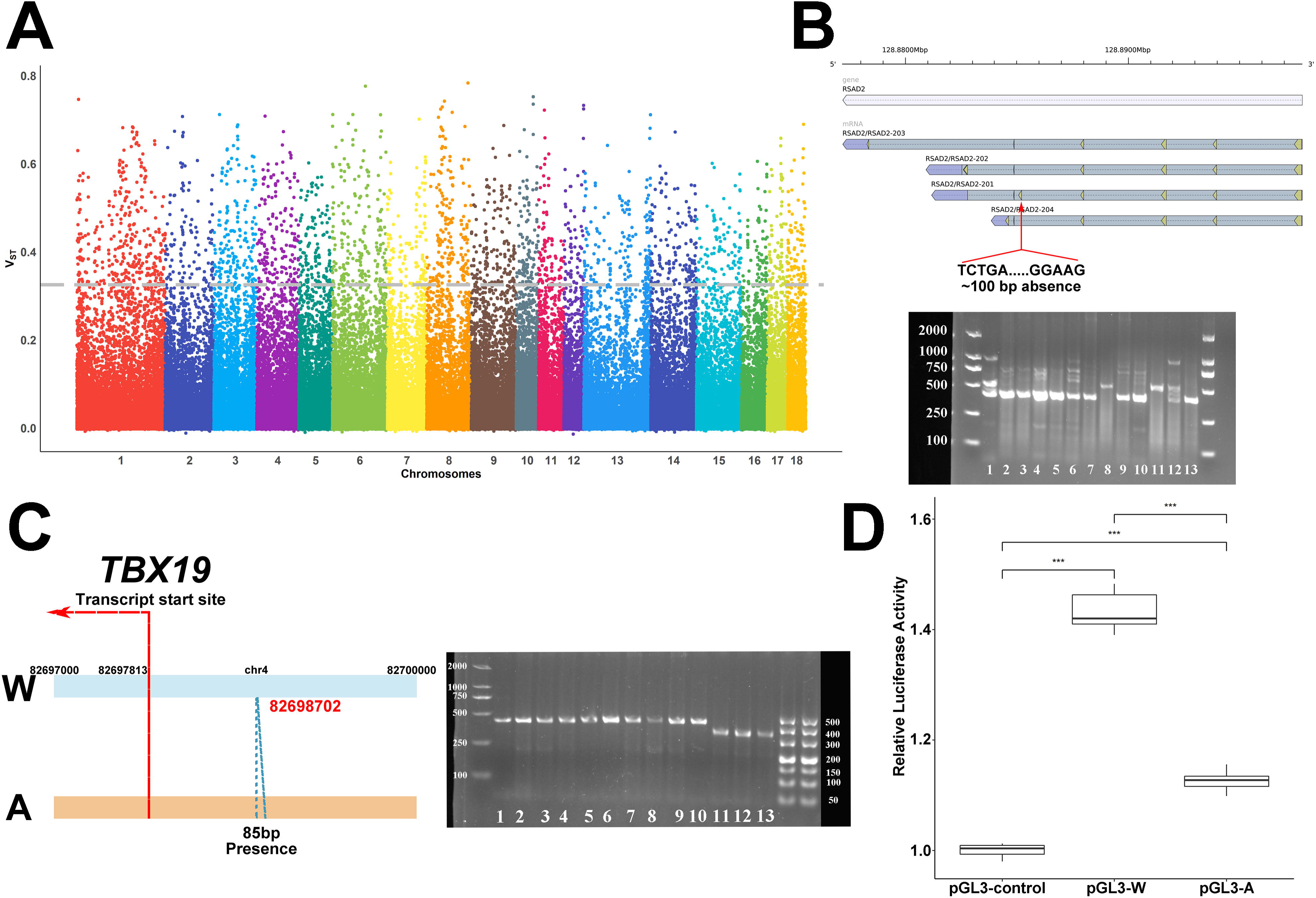
Genome-wide screening and selective signatures between European and Asian pigs. **A**. Statistic *V*_ST_ was plotted for selected SVs through pairwise comparison with threshold *V*_ST_ value ≥ 0.33. **B.** The gene structure of the *RSAD2* gene and the breakpoint of absence in its exonic region. The chart below indicates the PCR results for detecting this deletion in diverse breeds. The numbers from 1 to 13 represent the breeds as follows: Wuzhishan pig, Tongcheng pig, Bama Xiang pig, Rongchang pig, Ningxiang pig, Meishan pig, Diannan Small-ear pig, Bamei pig, Jinhua pig, Tibetan wild boar, Large White pig, Landrace and Duroc. **C.** Genomic structure of different alleles in the promoter region of *TBX19* gene. The blue lines between the two horizontal bars represent the presence of allele A. The right plot indicates the PCR results for detecting the presence of diverse breeds. The numbers from 1 to 13 represent the breeds in the same way as in Figure 6B. **D.** Comparison of transcriptional activity among different *TBX19* promoter regions in 293 T cell line.

Previous enrichment analyses revealed the enrichment of SVs within promoter regions; thus, we also endeavored to discover genes with promoters under selection. A total of 35 genes were identified, and nine were enriched in the cytosol Gene Ontology (GO) terms (Table S15). We also performed an SNP-based selective sweep analysis based on the same populations using the fixation index (*F*_ST_) and nucleotide diversities (π) method (Figure S12). Enrichment analysis suggested that the SVs under selection showed significant enrichment in SNP-based selective sweep regions compared to the random background model (P < 0.01, Figure S12). In total, 62 SVs were under potential selection within the significant selective sweep regions detected by the SNPs. More importantly, the SV in the promoter of *TBX19* was under significant selection based on both SV and SNP detection. To further dissect the function of this SV, we first analyzed the PCR results for this SV breakpoint and identified two alleles (Figure 6C), which were defined as W (wild type) and A (85bp insertion at the promoter of *TBX19*) alleles. To further verify the molecular effects of this insertion, luciferase expression levels were investigated to determine transcriptional activity by transfecting two types of recombinant plasmids (pGL3-A and pGL3-W) into 293T cells. Before performing the luciferase activity experiment, we screened the genome region that contained the pGL3 construct and confirmed that no differences occurred except for the A insertion. Therefore, the difference in activity between the two constructs was due to the insertion. The activities of the pGL3-A and pGL3-W groups were higher than those of the pGL3-control group (*t*-test). Moreover, the transcriptional activity of the insertion (A) was significantly lower than that of the wild type (W) (*t*-test, P = 3.73×10^-5^; Figure 6D). These results suggest that this insertion may regulate *TBX19* expression.

### The difference in the Y chromosome between European and Asian boars

Our study provides long-read sequencing reads from three male pigs and offers an opportunity to discover differences between European and Asian boars based on the Y chromosome. The Y chromosome of Sscrofa11.1 was used as the reference Y genome, and the long reads of four Chinese domestic pigs (Laiwu, Meishan, Tongcheng, and Bama Xiang) and a European hybrid pig (USMARC) were aligned to the reference. We finally detected 1485 SVs on the Y chromosome (**Figure 7A**). A total of 146 SVs were detected in all four Chinese domestic boars; however, not in the European hybrid porcine genome; 31 SVs were in the pseudoautosomal region (PAR), whereas the others resided in the non-pseudoautosomal region (NPAR), and 77.40% of the 146 SVs were in intergenic regions. In addition, we observed a new sequence inserted in the exon of the *LOC100624149* gene (Figure S13). This gene, also known as the *EIF2S3Y* gene, was previously reported to be associated with spermatogenesis [34]. In addition, in the NPAR region of the Y chromosome, we identified a new insertion in the intron of the *ZFY* gene (Figure 7B). Furthermore, we identified distinct haplotypes of this gene between European commercial pigs and Asian indigenous pigs (Figure 7C, File S1).

**Figure 7.**
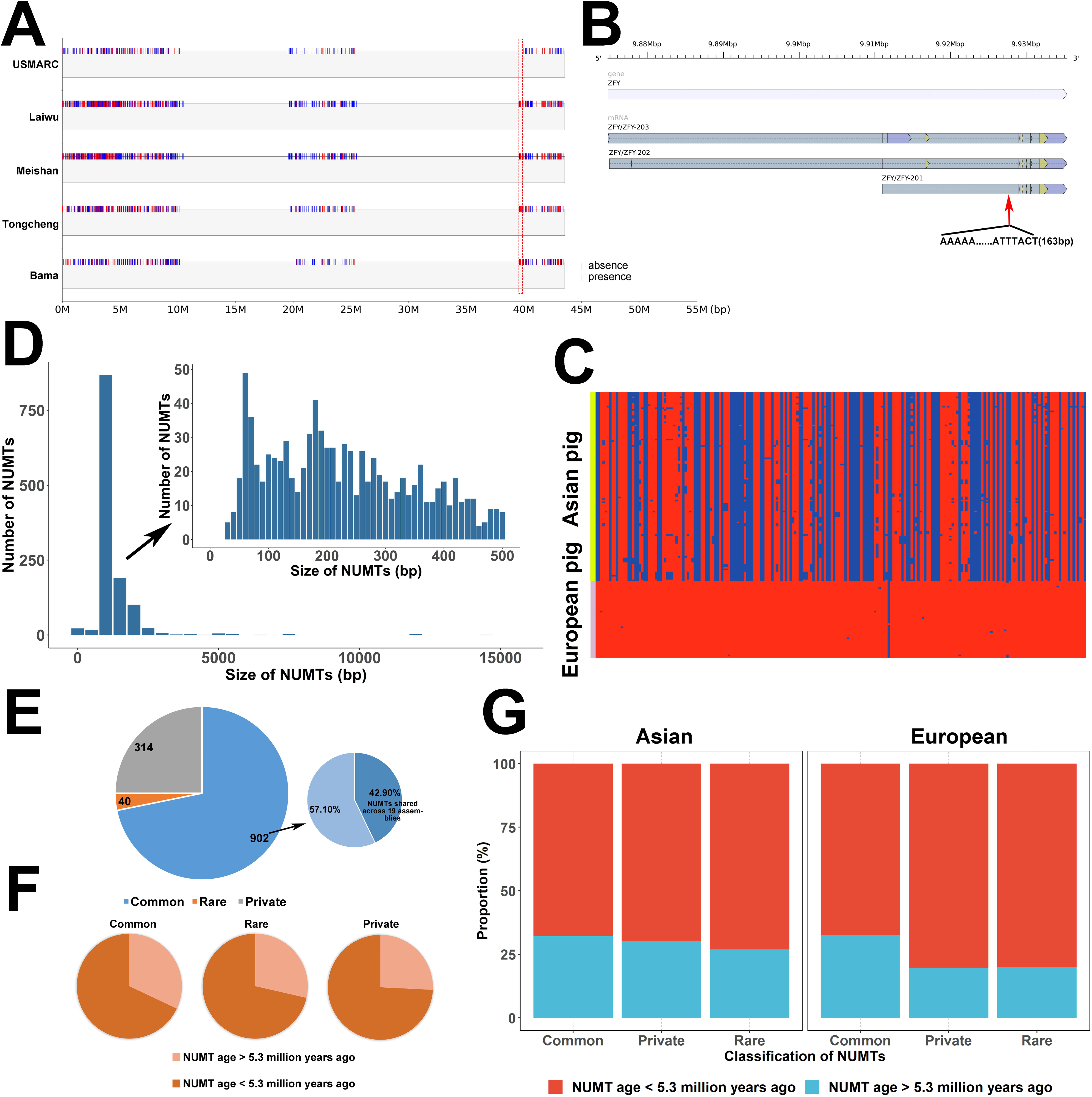
The SVs detected in the Y chromosome and the distribution of NUMTs in the porcine genome. **A**. The SVs detected in the Y chromosome using long reads from five boars. The red dashed rectangle indicated the regions where Asian pigs contained SVs while Europeans did not. **B.** The gene structure of the *ZFY* gene and the breakpoint of presence in its intronic region. **C.** The SNPs in the *ZFY* gene region across the Eurasian pigs. Blue blocks represented the existence of SNPs, while red blocks indicated not. **D.** Size distribution of NUMTs. NUMTs smaller than 500 bp were shown in the inset. **E.** The left pie plot indicates the proportion of three different NUMT categories. The right pie plot shows the proportion of NUMTs detecting in all 19 assemblies, or frequency > 0.1 in all assemblies. **F.** The proportion of older and younger NUMTs among common, rare, and private NUMTs. **G.** The proportion of NUMTs observed in older and younger NUMTs in European and Asian pigs, respectively.

In addition, in the Y chromosome, we noticed a 394 kb region (39.56 Mb – 39.95 Mb) containing SVs only found in Chinese domestic pigs (Figure 7A). Further analysis of this region compared to the other regions of the Y chromosome revealed that the SNP density of this region (0.04 variants/kb) was extremely low relative to the average density (0.59 variants/kb). More importantly, most SNPs in this region (72.73%) were rare variants (< 0.05) and might have little influence on this genomic region. Five of the 17 SVs in this region were detected in all four Chinese domestic boars, and an approximately 7 kb deletion was identified in the intronic region of the *LOC110257970* gene (Figure S14, Table S16). The *LOC110257970* gene only contained this SV, which was detected in the four Chinese domestic boars, and this deletion was adjacent to the 5′UTR (3.5 kb) of this gene. This gene is a BCL-6 corepressor-like gene, and one of its conserved domains is homologized to the non-ankyrin-repeat region of the *BCOR* gene, which may be related to germinal center formation and apoptosis [35].

### Characterization of the nuclear DNA of mitochondrial origin in the pig genome

We used a validated nuclear mitochondrial insertions (NUMTs) detection pipeline [36] to recognize the NUMTs in the 19 assemblies. In total, 12,002 NUMTs were identified, and a typical assembly carried a median number of 613 NUMTs, with sizes ranging from 34 to 15,975. After removing redundant NUMTs, we achieved 1256 NUMTs across the 19 genomes (Figure S15). Most NUMTs were short insertions; 50.80% of NUMTs were less than 300 bp, and 71.10% were less than 500 bp (Figure 7D). We cataloged the NUMT datasets into three classes (Figure 7E): common (frequency (*F*) ≥ 0.1), rare (*F* < 0.1), and private (detected only in one assembly). In total, 902 distinct NUMTs (71.82%) were common, including 387 shared across all the genomes, 40 rare NUMTs, and 314 private NUMTs. Three new genomes were assembled in this study that combined rare and private NUMTs with common NUMT data; thus, we identified 29 NUMTs that, to our knowledge, have not been reported previously. We compared the NUMTs to the annotation of the Sscrofa11.1 reference genome and identified 95 germline NUMTs located in gene regions, with the majority (82.11%, *n* = 78) being enriched in introns against exons (Table S17). The NUMTs located in the CDS of the reference were common (*F* ≥ 0.1) for the 19 assemblies, and these influenced genes were all novel genes.

To understand the molecular evolution of NUMTs, we estimated when these insertions occurred in the Suinae lineage by comparing modern mtDNA sequences with a consensus Suinae mtDNA sequence and detecting diagnostic sites that matched specific positions in each NUMT. Finally, we estimated the ages of the 657 NUMT insertions. The majority (69.71%) of NUMTs were less than 5.3 million years old, with 349 originating during the Pliocene period (2.5 – 5.3 million years ago). As expected, older NUMTs were more common in the population (Figure 7F). Notably, the private and rare NUMTs in European pigs were younger than those in Asian pigs (Figure 7G). Overall, our results indicate ongoing NUMT insertion and evolution throughout pig evolution.

## Discussion

Pigs are domesticated animals that play a vital role in providing sufficient protein resources to populations worldwide. Obtaining a more profound understanding of the legacy of latent genomic variation in porcine diversity will significantly benefit the development of animal husbandry and provide resources for the broader biological and genomic research communities. In this study, we performed *de novo* assembly and annotation of three new male porcine genomes by integrating data from PacBio sequencing, paired-end short-read sequencing, and Hi-C technologies. More importantly, we constructed a pangenome of the pig through assembly and comparative analyses of three new chromosomal-level genomes and 16 publicly available genomes representing diverse populations. Remarkably, our analyses detected a large number of genetic variants among the 18 genomes and Sscrofa11.1 reference assembly (an average of 10,215,593 SNPs, 71,704 SVs, and 68,538 PAVs) compared to previous SV studies based on short-read DNA sequencing data [37,38]. In addition, to limit contamination by new sequences, only non-reference sequences larger than 300 bp were retained from the previously published porcine pangenome based on 11 contig-level chromosomes; therefore, some variants might have been omitted. In this study, using the new chromosomal-level genomes, we detected a 66.17 Mb new sequence that complemented the previous pangenome. Furthermore, our enrichment analysis of SVs and genomic features revealed strong associations, particularly the frequent occurrence of absences around promoters or enhancers, thereby demonstrating the potential function of SVs in influencing gene expression. The newly constructed pangenome offers insights into genomic variation across Eurasian indigenous pigs and provides valuable tools for functional genomic analyses in pigs.

Indigenous Asian and European pigs are independently domesticated in Asia and Europe. Previous studies have reported that long-term independent domestication has left unique genomic footprints in Eurasian domestic pigs, such as selective sweeps of European variants around the *NR6A1*, *PLAG1*, and *LCORL* genes [39] and strong selective sweep signals at *GPR149* and *JMJD1C* on genes in Tongcheng pigs [40]. In this study, we focused on the distribution of SV hotspots in European and Asian pigs. We discovered only a long SV hotspot cluster region in chromosome X (49.80 – 100.50 Mb) of Asian pigs. This result corresponds to a previously reported X-linked selective sweep region in Eurasian pigs [5] and confirms the differences between European and Asian pigs in this region. In addition, we detected a new region that contained only SV hotspots in European pigs and located the important *CRPPA* gene, which is vital for skeletal muscle development, structure, and function [24]. To understand the contribution of SVs to the divergence between European and Asian pigs, we employed SVs to identify the regions under selection in Asian pigs compared with those in European pigs. The selection regions identified by SVs tended to overlap with the selective sweep regions detected by SNPs in our enrichment analysis, although with only a modest number of overlapping windows. This result is consistent with previous SV studies on rice [41]. In the overlapping regions, we identified an SV residing in the promoter of the *TBX19* gene. Previous studies have detected the *TBX19* gene under selection using SNPs [42,43]. *TBX19* is a protein-coding gene belonging to the T-box family of transcription factors, which play pivotal roles in regulating developmental processes [44]. Functionally, *TBX19* serves as a positive regulator of pro-opiomelanocortin (POMC) expression, thereby influencing the production of adrenocorticotropic hormone (ACTH) within the hypothalamic-pituitary-adrenal (HPA) axis. Deficiency in ACTH is associated with symptoms of adrenal insufficiency, including weight loss, reduced appetite, obesity, hypoglycemia, and low blood pressure [45]. Previous studies have reported that this gene is related to development, growth, and timidity traits in Chinese native pigs [42,43]. Our luciferase reporter assay further indicated that the newly detected SV in the promoter of the *TBX19* gene has the potential to alter the expression of this gene. Furthermore, H3K27ac enhancer research validated the high-activity peaks that existed within the promoter of the *TBX19* gene in Bama Xiang pig [46]. Overall, the SV identified under selection appeared to have a regulatory effect on the expression of the *TBX19* gene, making it a promising candidate for further exploration in studying developmental and timidity traits in Chinese domestic pigs.

In this study, our pangenome graph highlighted the importance of SVs in capturing missing heritability. In particular, with the addition of SVs, the estimated heritability significantly increased compared to that estimated using SNPs, consistent with a previous study [16]. In addition, we found that SVs contributed the largest share of heritability for ∼6.14% of molecular traits, which indicated that the detected SVs could be excellent genetic markers to compensate for the difficulty of elucidating traits by SNPs. Additionally, numerous studies have demonstrated that SVs can cause major phenotypic variations that affect a series of important traits [7,39] and may complement SNP-GWAS in identifying associations with phenotypes [47]. In the present study, we detected a lead SV that correlated with the abundance of *Bacteroidales* order. The lead SV was located in the intron of the *DSCAM* gene, which plays an important role in immune specificity and memory [28,48]. This SV and its genes could serve as candidate markers for discovering the relationship between the host and its microbiota. In conclusion, the pangenome graph and detected SVs provide new insights for studying complex porcine traits.

Our study offers an opportunity to investigate the divergence of Y chromosomes between European and Asian pigs in detail using long-read sequencing. We identified different haplotypes in the *ZFY* gene for European and Asian pigs. In humans, this gene regulates the transcription of some Y-linked genes [49] and mediates multiple aspects of spermatogenesis and reproduction, such as capacitation, acrosome reaction, and oocyte activation [50]. The haplotype disparity of this gene between European and Asian pigs revealed the diversification of Eurasian pigs, which might be related to discrepancies in reproductive traits. In addition, a 394 kb region where only Asian pigs contained SVs was discovered, and this region exhibited significantly lower SNP density than other regions. Hence, this intriguing region is easy to omit using SNPs. In this region, we detected an approximately 7 kb deletion in the intronic region and near the 5′UTR of the *LOC110257970* gene. This gene is a BCL-6 co-repressor-like gene homologized to the human *BCOR* gene. *BCOR* plays a critical role in early embryonic development and may also be involved in specifying the left and right sides of the body in developing embryos [35]. This discovery of unique SVs in Chinese domestic pigs may be a potential marker for studying the differences between European and Asian pigs during embryonic development.

## Materials and methods

### DNA extraction and genome sequencing

Genomic DNA was isolated using the DNeasy Blood & Tissue Kit (QIAGEN, Northrhine-Westfalia) and blood from three Meishan, Tongcheng, and Laiwu pigs. The integrity of the DNA was determined using an Agilent 4200 Bioanalyzer (Agilent Technologies, Santa Clara, California). Briefly, 8 mg of DNA was sheared using g-tubes (Covaris, Woburn, Massachusetts) and concentrated using AMPure PB magnetic beads (Pacific Biosciences, Menlo Park, California). Each SMRT bell library was constructed using a Pacific Biosciences SMRT bell Template Prep Kit (Pacific Biosciences, Menlo Park, California). The constructed libraries were size-selected on a BluePippin system (Sage Science, Beverly, Massachusetts) to isolate molecules of ∼ 20 kb, and a Sequel Binding and Internal Control Kit 3.0 (Pacific Biosciences, Menlo Park, California) was used for primer annealing and binding of the SMRT bell templates to the polymerase process. Sequencing was performed on a Pacific Biosciences Sequel II platform (Pacific Biosciences, Menlo Park, California) using 18 SMRT cells.

### Hi-C library construction and sequencing

Approximately 10 mL of blood drawn from each sample was used for the Hi-C experiment. Blood was first crosslinked in a 2% formaldehyde solution for 15 min, and the crosslinking reaction was stopped by adding glycine. After the nuclei were isolated, the chromatin was digested with *MboI*. The sticky ends of the digested fragments were biotinylated, diluted, and ligated randomly. The DNA fragments labeled with biotin were sheared by ultrasound, blunt-end repaired, and A-tailed. The adapters were then ligated to the DNA fragments, and PCR amplification was used to construct the Hi-C library. After quality control, the Hi-C library was sequenced using an Illumina paired-end sequencing platform with 2×150 bp reads.

### Transcriptome sequencing

Transcriptome sequencing was performed on 37 tissue samples from the liver, spleen, kidney, lung, thymus gland, stomach, duodenum, lymph, ovary, and muscle isolated from three individuals. Total RNA was extracted from each tissue sample using a TRIzol-based RNA extraction kit (Invitrogen, Carlsbad, California). RNA degradation and contamination were monitored using 1% agarose gel electrophoresis. The concentration of total RNA was measured using a Qubit RNA Assay Kit on a Qubit 2.0 Fluorometer (Life Technologies, Carlsbad, California). RNA sequencing libraries with 250–350 bp insert sizes were prepared using Kapa RiboErase (Roche, Basel, Switzerland). All libraries were sequenced on the Illumina NovaSeq 6000 S4 platform according to the manufacturer’s instructions to obtain transcriptome profiles.

### Genome assembly

All subreads from PacBio sequencing for each breed were assembled using Falcon (v2018.03.12) [51]. The assembled genomes were then polished using Pilon (v1.23) [52] with the filtered Illumina paired-end reads described above. Several rounds of iterative error correction were performed to ensure the accuracy of the genomes. Reads from the Hi-C library were then used to construct a pseudo-chromosome. After removing the adapter sequences and low-quality bases, these reads were aligned to the corresponding assembly using the aln and sample commands from bwa (v0.7.17). The resulting BAM files and contigs from the assembly were used as inputs for LACHESIS (https://github.com/shendurelab/LACHESIS) with the cluster number set to 20 and anchored to the pseudo-chromosome. Finally, chromosome-level genomes were manually optimized using JuiceBox (v2.20.00) [53].

### Genome assembly assessment and annotation

Our assembled genomes and the other 16 public genomes used in this study were assessed using BUSCO (v5.0.0) [21] based on the lineage dataset vertebrata_odb10 (creation date: 2019-11-20). RepeatMasker (v4.1.2) (http://www.repeatmasker.org) was used to detect repeats in genomes. Gene prediction was conducted by combining three independent approaches in each repeat-masked genome, including *ab initio* prediction, homology-based prediction, and transcriptome-based prediction. For *ab initio* gene prediction, BRAKER2 (v2.1.6)[54] and GlimmerHMM(v3.0.4) were used with default parameters. For homology-based prediction, protein sequences from humans (*Homo sapiens*), mice (*Mus musculus*), cows (*Bos taurus*), sheep (*Ovis aries*), and Sscrofa11.1 reference pig genome (*Sus scrofa*) were supplied, and the gene models were predicted by GeMoMa (v1.9)[55]. For transcriptome-based prediction, the RNA-Seq data of each breed were aligned to the corresponding assembly by HISAT2 (v2.2.1) with default parameters. StringTie (v2.1.6) and TransDecoder (v5.5.0, https://github.com/TransDecoder/TransDecoder) were then used to assemble the transcripts and convert candidate coding regions into gene models. Simultaneously, these RNA-Seq data were also *de novo* assembled by Trinity (v2.1.1), and PASA (v2.5.3) was utilized to predict the gene structure. Finally, the gene models predicted through the above three approaches were combined by EvidenceModeler (v2.1.0) [56] into a non-redundant set of gene structures.

### Core and dispensable gene family clustering

The core and dispensable gene sets were estimated based on gene family clustering using OrthoFinder (v2.5.4) [57]. For each genome, a gene whose CDS was 100% similar to that of the other genes was removed using the cd-hit-est function of CD-HIT (v4.8.1) [58]. The protein sequences of the remaining genes were processed using OrthoFinder with diamond (v2.0.7.145). Based on the results, gene families were categorized into four classes: core, soft core, dispensable, and private gene families. We used InterProScan (v5.47-82.0) [59] to annotate the protein domain using Pfam and GO datasets.

### SNP analysis of different breeds

The remaining 18 genomes were aligned to the reference genome Sscrofa11.1 using Mummer (v4.0.0rc1) [60] with the parameter settings “-g 1000 –c 90 –l 40.” The alignment block was then filtered out of the mapping noise, and the one-to-one alignment was identified using a delta-filter with the parameter setting “-1.” Show-snps was used to identify SNPs with the parameter setting “-ClrT.” All clean reads for the 599 pigs across Eurasian were mapped to the Sscrofa11.1 genome using the “MEM” function of bwa. SAMtools (v1.15) was used to sort the mapped reads, and samblaster (v.0.1.26) was applied to mark potential PCR duplications. The GATK (4.1.2.0) HaplotypeCaller best practice was used to detect SNPs. Obtained SNPs were filtered using the VariationFiltration in GATK, according to the following criteria: (1) approximate read depth >10×; (2) variant confidence/quality by depth >2.0; (3) RMS mapping quality (MQ) >40.0; (4) Phred-scaled P value using Fisher’s exact test to detect strand bias <60.0; (5) Z-score from the Wilcoxon rank sum test of Alt vs. Ref read MQs (MQRankSum) >−12.5; and (6) Z-score from the Wilcoxon rank sum test of Alt vs. Ref read position bias(ReadPosRankSum) >−8.0.

### Structural variation identification

Previously filtered Mummer alignment delta files were used to detect structural variations using smartie-sv (https://github.com/zeeev/smartie-sv) with default parameters. The detected variations from smartie-sv consisted of insertions and deletions; meanwhile, insertions/deletions were treated as the PAV region. Simultaneously, the SyRI pipeline [61] was applied based on filtered Mummer delta files with default parameters to detect inversions and translocations. According to the definitions of sequence variation in SyRI outputs, we converted the INV variants into inversion SVs relative to Sscrofa11.1. Simultaneously, TRANS and INVTR were considered translocation SVs relative to Sscrofa11.1.

### Identification of SV hotspot regions

We calculated the distribution of SV breakpoints for each 200 kb window (with a 100 kb step size) along each chromosome. Then, all 200 kb windows were ranked in descending order according to the number of SVs within the window. The top 5% of all windows with the highest frequency of SV breakpoints were regarded as SV hotspots. All consecutive hotspot windows were merged as the “hotspot regions.” Simultaneously, we employed the same method to identify the hotspot regions for each subpopulation.

### Graph-based pangenome construction and utilization

SVs from the 18 genomes were filtered using a previously reported pangenome construction pipeline [9]. In brief, redundant SVs were removed, and the remaining SVs with more than 90% repetitive sequences were omitted. The Sscrofa11.1 genome was set as a reference and combined with the non-redundant SVs to build the graph pangenome using vg (v1.56.0) [62]. The “Giraffe” pipeline was used for SV genotyping [63], and the resulting variant call format file for each individual was merged using VCFtools (v0.1.17) [64].

The heritability of ASVs and gene expression was estimated using a previously reported pipeline [16]. Briefly, the following genetic variants were removed: (1) mean depth < 5×, (2) missing rate > 95%, and (3) minor allele frequency < 0.1. For SNPs, we performed LD-based pruning using PLINK (v.1.9) [65] with the “--indep-pairwise 50 10 0.1” option. While, the LD-based pruning was not performed on SVs. The LDAK-thin model [27] was applied to estimate the proportion of phenotypic variance explained by genetic variants. Finally, we added the parameter “--constraint YES” to ensure that all estimations with LDAK-thin are bounded within [0,1].

We performed GWAS using a mixed linear model (MLM) method. Furthermore, we used the leave-one-chromosome-out (LOCO) method and association studies implemented in GCTA (v1.93.2beta) [66]. Genetic variants in the heritability analysis were used in the genetic relationship matrix (GRM). A constant threshold of 1×10^-6^ was used as the significance threshold. The eQTLs analysis was performed by MatrixEQTL (v2.3).

### Detection of selective sweeps

For SVs, we tested the selective sweeps between European commercial pigs and Asian indigenous pigs using the *V*_ST_ approach. *V*_ST_ [33], a static analog of *F*_ST_, estimates population differentiation based on quantitative intensity data and varies from 0 to 1. The statistical *V*_ST_ of each SV was calculated to determine the selective signals between the European and Asian domestic pigs. *V*_ST_ was calculated as follows:

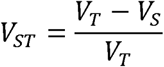

where *V*_T_ is the variance of SVs among all unrelated individuals in the target and control populations, and *V*_S_ is the average variance within each population, which was weighted for population size. SVs with the top 1% *V*_ST_ values were considered significantly under selection.

We scanned the selective signals for SNPs using *F*_ST_ and nucleotide diversity (π) values. The *F*_ST_ and π values were calculated using VCFtools with sliding windows, performed with a 100 kb window size and 10 kb step size. Selective sweeps identified by SNPs were determined by merging continuous genomic regions displaying the top 5% *F*_ST_ and π (European pigs/Asian pigs) values.

### Paternal analysis and NUMT detection

We aligned the long reads of five males to the Sscrofa11.1 reference genome using minimap2 (v 2.17) and applied sniffles (v 2.0.7) to detect SVs on chromosome Y. The NUMTs in the 19 assemblies were detected using a previously reported method [36]. In brief, two copies of the mitochondrial sequence in Sscrofa11.1 (AF486858.1) were concatenated and then compared to the genome assemblies using GABLAM [67], which is a wrapper for BLAST+ to identify potential NUMTs. The NUMT age was estimated using a previously described method [68]. We aligned the mitochondrial sequences from Sscrofa11.1 for the common warthog (*Phacochoerus africanus*, DQ409327.1) and the consensus sequence from each NUMT contig using MUSCLE (v3.8.31) [69]. We counted the number of sites within each NUMT region where modern and ancestral mitochondrial sequences differed. The ratio of sites that matched the modern allele to all different sites was calculated and used to derive an approximate age for each NUMT relative to the estimated *Sus scrofa* – *Phacochoerus africanus* divergence time of 9.7 million years [70]. To ensure the accuracy of the results, we only retained NUMTs with lengths between 50 and 1,000 bp and excluded NUMTs without different alleles between Sscrofa11.1 and the common warthog.

### SV validation

The SVs under selection located in the CDS and promoter regions were validated using specific PCR assays for the 13 distinct Eurasian pig breeds. Primers were designed for each SV using Primer3 (v4.0.1) based on sequences around the breakpoints in Sscrofa11.1. The PCR amplifications were performed in 25 μL reaction volumes using the 2×EasyTaq PCR SuperMix (Trans, Beijing) and processed as follows: initial denaturation at 94 L for 5 min; amplification for 32 cycles at 94 L for 30 s, 58 L for 30 s, and 72 L for 1 min; and a final extension at 72 L for 5 min.

### Luciferase reporter assay of the *TBX19* promoter region

The full-length promoter of *TBX19*, with or without the insertion, was synthesized separately by chemical synthesis. Then, the two types of *TBX19* promoter regions were cloned into the pGL3-Basic luciferase vector (Jinkairui, Wuhan) using the NheI/XhoI restriction sites (Jinkairui, Wuhan). Sequencing was performed to verify the plasmid identities. All recombinant plasmids, the pRL-TK plasmid (Jinkairui, Wuhan) and Lipofectamine 2000 (Invitrogen, Carlsbad, California), were transfected into the 293 T cell line. After 48 h, firefly and Renilla luciferase activities were measured and investigated using the Dual-Luciferase Reporter Assay System (Yeasen Biotechnology, Shanghai).

## Ethical statement

This study was conducted in strict accordance with the protocol approved by the Institutional Animal Care and Use Committee (IACUC) at the China Agricultural University (Beijing, People’s Republic of China; permit no. DK996).

## Data availability

Assembled genomes and raw sequencing data that support the findings of this study are publicly accessible and have been deposited into NGDC database (https://ngdc.cncb.ac.cn/) with accession number PRJCA019609. The genome annotation results, graph-based pangenome, SNPs, and SVs detected in this study can be accessed by Zenodo (https://doi.org/10.5281/zenodo.12794748) or NGDC under accession code GVM000861 (https://ngdc.cncb.ac.cn/gvm/getProjectDetail?project=GVM000861).

## Code availability

The codes used in the study are available from the GitHub repository (https://github.com/kimi-du-bio/GBPPG).

## CRediT author statement

**Jian-Feng Liu:** Conceptualization, Methodology, Supervision, Writing – review & editing. **Heng Du:** Investigation, Writing – original draft. **Yue Zhuo:** Investigation, Writing – original draft. **Shiyu Lu:** Validation. **Wanying Li:** Validation, Writing – review & editing. **Lei Zhou:** Writing – review & editing. **Feizhou Sun:** Resources, Writing – review & editing. **Gang Liu:** Resources.

## Competing interest

The authors have declared no competing interests.

## Acknowledgments

This work was financially supported by the Biological Breeding-Major Projects in National Science and Technology (2023ZD0404405), the National Natural Science Foundations of China (32272844 and 32302708), Chinese Universities Scientific Fund (2023TC196), Science and Technology Program of Guizhou Province (Qian Kehe Support [2022] Key 032), the Earmarked Fund for China Agriculture Research System (No. CARS-pig-35) and the 2115 Talent Development Program of China Agricultural University. We would like to thank High-performance Computing Platform of China Agricultural University for computing supporting.

## Supplementary materials

### Supplementary files

**File S1. Supplementary information**.

### Supplementary figure legends

**Figure S1. The neighbor-joining tree was constructed by 599 genomes.**

**Figure S2. The frequency distribution of *k*-mer for each genome (*k*=17)**.

A. Laiwu pig. **B.** Meishan pig. **C.** Tongcheng pig.

**Figure S3. Genome-wide contact matrix of three pig genomes. The color intensity represents the frequency of contact between two 500 kb loci**.

**B.** Tongcheng pig. **B.** Laiwu pig. **C.** Meishan pig.

**Figure S4. The SNPs were identified by assembly-comparison for each assembly.**

**Figure S5. Correlation of SNP density, π, Ts/Tv, and Tajima’d from 18 *de novo* assembled genomes (compared to the reference genome) and 599 resequencing samples**.

**A**. SNP density (number of variants in 1kb region). **B.** π. **C.** Ts/Tv. **D.** Tajima’d.

**Figure S6. Characterization of SVs in the 18 pig genomes. The heatmap shows the present frequency of SVs in 18 genomes**.

**A.** Absence. **B.** Presence. **C.** Inversion. **D.** Translocation.

**Figure S7. Chromosome distributions of hotspot SVs in European pig population, BL, DWB, RT, and TNJM populations.** The red box indicates the region with hotspots that have not been detected in the European population, while the blue box indicates that only the European population detects the hotspots.

**Figure S8. The annotation results of PAVs with genes and their flanking regions.**

**Figure S9. The pangenome graph shows variations within 2175210-2175820 bp on chromosome 1.**

**Figure S10. The GWAS result of ASV2205**.

**A.** The Manhattan plot. **B.** The quantile-quantile plot.

**Figure S11. The RNA-Seq alignment results of *RSAD2***.

**Figure S12. The genomic regions under selection are inferred by SNPs**.

**A.** *F*_ST_ results. **B.** Nucleotide diversity result. **C.** The SNPs are selected by two methods; the red dot represents the select regions. **D.** The enrichment analysis of SVs under selection and the regions under selection detected by SNPs.

**Figure S13. The gene structure of the *LOC100624149* gene and the insertion in its exonic region**.

The red arrow represents the insertion position.

**Figure S14. The gene structure of the *LOC110257970* gene and the deletion in its intronic region**.

The blue rectangle represents the deletion region.

**Figure S15. The NUMTs positions are distributed in all chromosomes.**

### Supplementary tables

**Table S1. The 599 high-throughput Sequencing data used in this study**.

**Table S2. The pig genomes were collected from the previous studies.**

**Table S3. The detailed information for Illumina reads, PacBio reads, and Hi-C reads.**

**Table S4. Statistics of assembled contigs for three pig genomes.**

**Table S5. Profile of the annotated genes in the three new genomes.**

**Table S6. Evaluation of protein completeness for three new assemblies using BUSCO.**

**Table S7. Statistics profile of gene function annotation for new three genomes.**

**Table S8. The RNA-Seq data of 171 samples were used in this study.**

**Table S9. The 495 hotspots and related QTLs. Table S10. ASVs assigned taxonomy.**

**Table S11. The SVs have the high LD with the tag SNPs.**

**Table S12. The enriched KEGG pathways for genes are influenced by SVs under significant selection.**

**Table S13. 1005 QTLs overlapped with the genes influenced by SVs.**

**Table S14. Nine genes in which CDS was influenced by SVs.**

**Table S15. The enriched GO terms for genes whose promoters were influenced by SVs under significant selection.**

**Table S16. The five SVs in the 394 kb region of chrY and their overlapped genes.**

**Table S17. The genes overlapped with NUMTs.**

**Table S18. The primers used in this study.**

